# Early-warning surveillance of West Nile and Usutu viruses in water and mosquito excreta using digital PCR

**DOI:** 10.64898/2026.02.01.703130

**Authors:** Julien Mocq, Julie Raymond, Karine Bolloré, Anaïs Fossot, Rachel Beaubaton, Alexandre Lepeule, Cécile Gruet, Franz Durandet, Jeanne Hanin, Guillaume Lacour, Albin Fontaine, Olivier Courot, Antoine Mignotte, Yannick Simonin

## Abstract

West Nile virus (WNV) and Usutu virus (USUV) are mosquito-borne pathogens maintained in bird–mosquito cycles and increasingly cause human and equine disease in temperate regions. As most infections are asymptomatic, surveillance based on clinical and veterinary reports provides delayed signals for vector control and equine vaccination strategies. We implemented, in 2024 and 2025, an operational environmental surveillance strategy combining molecular xenomonitoring of adult mosquito excreta with water sampling from wetlands and mosquito breeding sites, coupled to a standardized multiplex reverse transcription digital PCR workflow. Viral detection was performed using a multiplex digital PCR assay with characterized limits of blank, detection, and quantification, and low-level positives were confirmed by Sanger sequencing. This multi-matrix approach detected WNV RNA in southern France as early as 1 July 2024, 40 days before the first human case and 67 days before the first equine alert, and on 9 April 2025, a full 19 weeks ahead of the first human case that year. USUV RNA was similarly detected from 5 July 2024 and 22 April 2025, providing actionable early warnings to guide targeted vector control and preparedness. Across both years, WNV and USUV were detected 29 and 8 times among 396 samples in 2024, and 90 and 14 times among 815 samples in 2025, revealing previously unrecognized, cryptic virus circulation. These results demonstrate that integrating environmental surveillance provides a sensitive, proactive framework for the early detection of emerging Culex-borne arboviruses, offering precious lead time for public one-health strategies.

## 2. Introduction

West Nile virus (WNV) and Usutu virus (USUV) are emerging threats in temperate regions, as climate warming, urbanization and biodiversity loss collectively alter the ecology, abundance and behaviour of mosquito vectors and avian reservoir hosts, promoting viral geographic expansion and increasing outbreak frequency (Brugger & Rubel, 2009; Erazo et al., 2024; Fay et al., 2025). These neurotropic orthoflaviviruses, members of the Japanese encephalitis virus serocomplex (Postler et al., 2023), are primarily transmitted by ornithophilic mosquitoes of the genus Culex (Clé et al., 2019; Goddard et al., 2002; Vogels et al., 2017), predominantly Culex pipiens in Europe (Vogels, et al., 2016; Zannoli & Sambri, 2019). A wide range of avian hosts serve as reservoirs for both viruses (Agliani et al., 2023; Constant et al., 2020), but non-avian vertebrates may be infected through infectious mosquito bloodmeals. In humans and horses, infections typically result in low viremia, making them epidemiological incidental “dead-end” hosts unable to infect subsequent biting mosquitoes (Gould et al., 2025; Kramer et al., 2008). Approximately 80% of WNV-infected horses or humans are asymptomatic, but clinical manifestations range from influenza-like symptoms to severe neuroinvasive diseases (Sejvar, 2007). USUV, usually causing mild or no symptoms, can trigger serious neurological complications, particularly in immunocompromised individuals (Cadar & Simonin, 2022; Vazquez et al., 2011). To date, no antiviral treatments or vaccines are available for WNV or USUV in humans (Vilibic-Cavlek et al., 2019).

Since the 1990s, recurrent WNV-epidemics of occurred in southern and eastern Europe (Zannoli & Sambri, 2019) while USUV remained relatively unnoticed until its emergence in 1996 (Engel et al., 2016) and spread across Europe in the early 2000s (Cadar & Simonin, 2022). In France, the first human and equine WNV cases were identified in 1962 in the Camargue region (Giudice et al., 2004), and the first human USUV case, in 2016 in Montpellier (Simonin et al., 2018). Since 2022, both Orthoflaviviruses expanded northward and westward (Chevalier et al., 2025; Duvignaud et al., 2024; Migné et al., 2024).The past few years, alongside multiple humans, equine, avian and mosquito USUV detections (Beaubaton et al., 2025; Bouchez-Zacria et al., 2025; Gonzalez et al., 2025), France experienced major WNV outbreaks of autochthonous human and equine cases, mainly in historically affected regions but also for the first time in more northern regions (Hassold-Rugolino et al., 2025). These patterns confirm active viral circulation and ongoing expansion beyond historical endemic areas in France and in Europe (Branda et al., 2025; Kirby et al., 2025).

The surveillance and control of WNV and USUV viruses in France remain limited by structural, financial, and operational constraints, impeding early detection and rapid intervention. Unlike for Aedes-borne viruses chikungunya, dengue, and Zika, France lacks a comprehensive risk-prevention and management framework for Culex-transmitted WNV and USUV. As a notifiable disease at the EU level, WNV surveillance typically begins after the declaration of a confirmed equine or human case of infection. Similarly, incidental USUV antibody detections during WNV serological screening suggest broader circulation than currently recognized (Clé et al., 2019; Simonin et al., 2018). Emergency adulticide operations are inadequate for WNV and USUV control, because of the long-range dispersal of Culex, and preventive larvicide application poses significant financial and operational challenges. These economic and logistical constraints, combined with knowledge gaps in early viral detection, highlight the limitations of current surveillance and vector control strategies for WNV and USUV. Molecular Xenomonitoring (MX) offers a non-invasive, cost-effective, and integrative alternative to the labour-intensive mosquito pooling and dissection, addressing the urgent need for timely arbovirus surveillance. It relies on detection of viral particles in mosquito excreta (Fontaine et al., 2016), subsequently analysed by molecular biology techniques. Viral RNA is only detectable in effectively infectious mosquitoes with disseminated infection, with RNA in head, malpighian tubules or both (Fontaine et al., 2016).

Moreover, the analysis of environmental matrices, particularly water, is a promising strategy of early detection of arboviruses WNV and USUV circulation in avian and mosquito compartments (Beaubaton et al., 2025; Kuhn et al., 2025). Virus particles in water can originated from oviposition fluids excreted by infected mosquitoes, which can potentially infect larvae by vertical transmission (Hamel et al., 2024a), dead infected mosquitoes, infected larvae or faeces of infected birds. Indeed, experimental studies found the genomes of WNV and USUV in the digestive systems of infected birds, subsequently excreted into the environment in the faeces (Beaubaton et al., 2025; Kipp et al., 2006), between three and eight days post-infection, peaking at day 5 (Kipp et al., 2006). While the presence of viral genomes does not necessarily imply the presence of an infectious virus, its detection in environmental matrices may serve as an early warning signal to trigger reinforced surveillance and vector control operations.

To improve early detection of WNV and USUV, this study evaluated an innovative approach combining extensive sampling with molecular analysis of mosquito excreta and water from potential larval breeding sites and wetland used as avian resting sites. We hypothesized that viral genomes could first be detected in water – introduced by infected bird excretion–, then subsequently appearing in local mosquito populations, ahead of equine or human cases. In 2024, field evaluations in areas of known or suspected viral circulation were conducted to optimize detection systems and establish proof of concept. In 2025, the method was deployed regionally to assess its performance and relevance for arboviral surveillance. We developed and validated a highly sensitive multiplex digital PCR assay for WNV, USUV, and mosquito-derived nucleic acids, together with optimized extraction protocols for both MX and environmental water samples. Over the two-year study, WNV and USUV were repeatedly detected across multiple matrices, often weeks before the first human or equine cases, with WNV lineage 2 confirmed by sequencing. These results demonstrate that integrating mosquito and environmental surveillance provides a sensitive, proactive framework for early detection of emerging Culex-borne arboviruses, offering unprecedented lead time to guide public health interventions in Europe.

## 3. Methods

### 3.1 Investigated matrices

#### 3.1.1 Mosquito excreta

The MX method was used to assess the circulation of WNV and USUV by detecting viral RNA traces in mosquito excreta, adapted and optimized for field surveillance (Fig.1A). Mosquitoes were trapped using BG-Pro or BG-Sentinel traps (Biogents, Germany) fitted with 3D-printed MX adapters (Bigeard et al., 2024). The conical collection nets of the traps were modified with elastic closures to ensure airtight contact with the MX adapter. Inside each adapter, a large feeder (30 ml capacity) with a cotton ball soaked in 10% honey water-maintained humidity and provided sustenance, enabling mosquitoes to survive during the collection period. At the bottom of the adapter, a 10 µm-thick aluminium foil ring (shiny side up) was placed to collect excreta and facilitate its recovery. After collection, mosquitoes were released in cage and sucked using a Prokopack (Vazquez-Prokopec et al., 2009), frozen at −20 °C, sexed, and identified to the genus level. During the 2024 field campaign, aluminium rings containing mosquito excreta (thereafter MX samples) were placed in plastic bags. For optimization purposes during the 2025 campaign, they were instead placed in 50 mL Falcon tubes designated for the elution step and subsequently stored at −80 °C until further processing for RT-dPCR analysis by IAGE (Ingénierie et Analyse en Génétique Environnementale, Montpellier, France).

**Figure 1:**
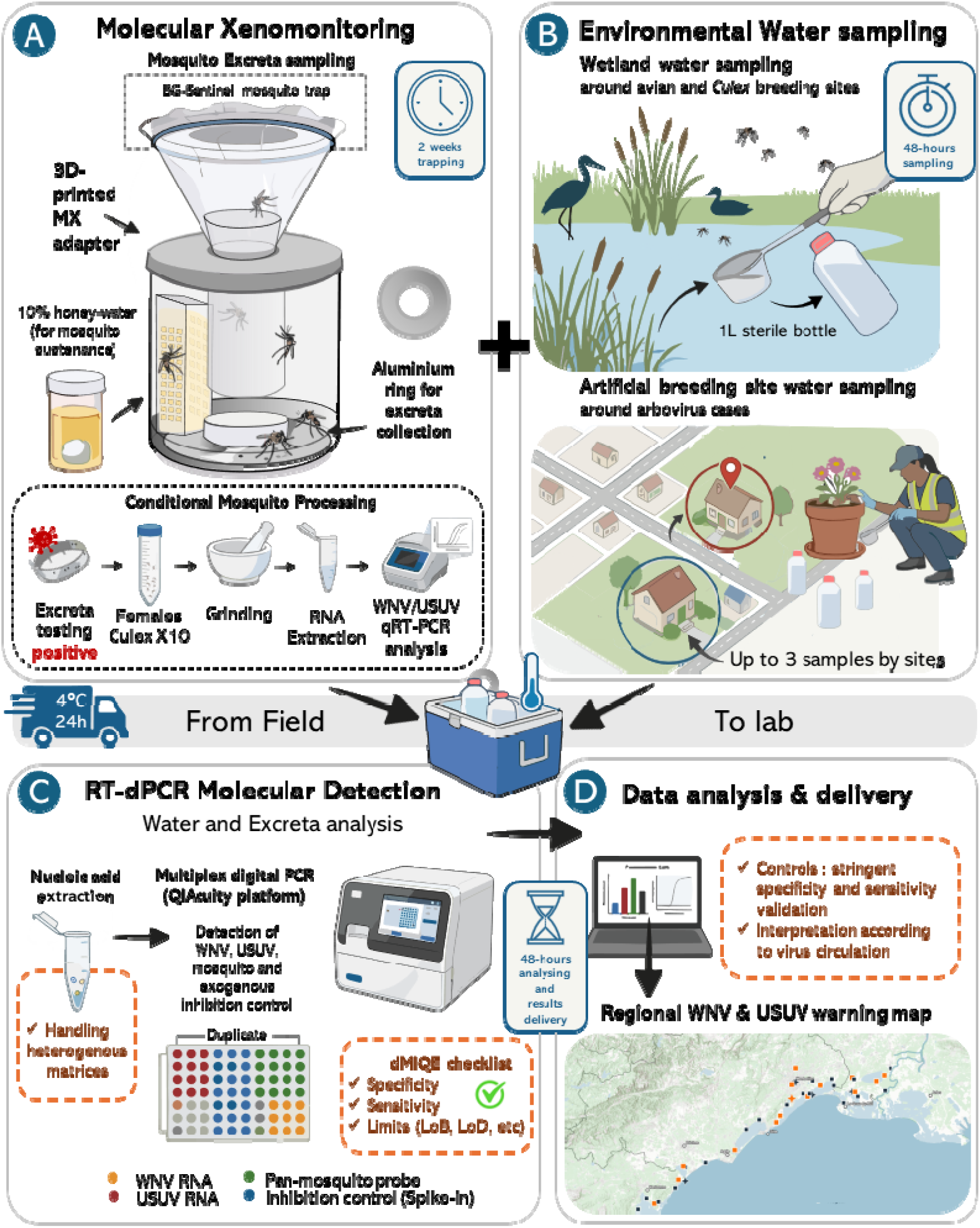
Early warning environmental surveillance workflow for WNV and USUV in 2024 and 2025 in Occitania, France.

Following a positive detection in excreta associated with the trapping, all Culex females from the positive trap were grinded by pool of 10 individuals, and viral RNA was extracted following Klitting et al.(Kittling & et al., 2026). Then, extracts were tested by WNV/USUV duplex qRT-PCR(Tinto et al., 2022).

#### 3.1.2 Wetland and artificial breeding site water collection

Environmental water sampling was carried out to detect viral RNA traces in wetland and breeding site habitats potentially harbouring infected avian and mosquito populations (Fig.1B). To maximise the chances of virus detection in wetlands, the survey prioritized ecologically and epidemiologically relevant environments for potential viral introduction and circulation: avian biodiversity hotspots and known bird congregation sites (Jourdain et al., 2007), such as zoological parks (e.g., Montpellier zoo (Beaubaton et al., 2025)); wetlands that serve as both nesting and migratory stopover zones for wild birds, and favourable habitats for Culex mosquitoes(Eiden et al., 2018; L’Ambert et al., 2012); sites with a documented history of WNV or USUV circulation, confirmed human or animal cases, or prior detections in mosquitoes in Occitania (Beaubaton et al., 2025; Murgue et al., 2001; Rodríguez-Valencia et al., 2025). In addition, in autumn 2024 during a significant circulation of WNV in equine population detected in Occitania(Grard et al., 2025), surveillance efforts were strategically intensified around equestrian facilities. These facilities are often located within or adjacent to wetland ecosystems, such as the Camargue Regional Natural Park and the Mediterranean coastal marshes, where domestic horses coexist with resident and migratory bird populations and dense Culex populations, creating key ecological interfaces conducive to arbovirus transmission. To ensure representative sampling and increase chances of viral detection, water from wetlands was collected in sterilized 1-liter polypropylene bottles from up to three points within a five-meter radius, by direct immersion of the bottles, or using a dipper, gently mixing the water column prior collection to homogenate surface and lower layers while avoiding sediment disturbance.

Waters from artificial breeding sites were mainly collected in urban areas by the mandated vector control operator Altopictus, in response to confirmed local arbovirus cases. Targeted entomological investigations, coordinated with the Regional Health Agency in Occitania (ARS Occitanie), involved door-to-door inspections aimed at identifying and eliminating water collections, potential and active mosquito larval breeding sites, colonized by Culex pipiens and Aedes albopictus. These interventions provided opportunities to collect water samplings in areas of attested arboviral circulation, in sterilized 1-liter polypropylene bottles. Small movable recipients (e.g. cups, vases…) were entirely drained, while unmovable breeding sites (e.g. neglected swimming-pool bottom) and nearby wetlands of interests were sampled with a mosquito dipper or a bulb syringe. Samples were kept at 4 °C during transport and delivered at the IAGE laboratory within 48 hours for molecular analysis.

### 3.2 Field protocols

#### 3.2.1 2024 operational proof of concept

Sampling was conducted from the 15^th^ of April to 23^rd^ of October 2024, corresponding to the active season of Culex spp. in Southern France (Lacour, 2016; Montarsi et al., 2015). Based on predefined ecological and epidemiological criteria, a total of 48 sampling sites were selected, across 13 municipalities within the Gard and Hérault departments (Fig.2A; Supplementary table S.1). Additionally, 12 municipalities were opportunistically sampled during vector control operations. Most traps operated autonomously using high-capacity batteries (27,000 mAh) allowing 3 to 4 days of function, with weekly or biweekly changing for repeatedly sampled sites; a few traps were exceptionally powered by mains electricity at fixed or secured locations. Two attractants were used: an olfactory lure BG-Mozzibait and CO₂ from BG-CO_2_ Generators (Biogents, Germany), both effective for attracting Culicidae species (Cilek et al., 2024; Wilke et al., 2019).

**Figure 2:**
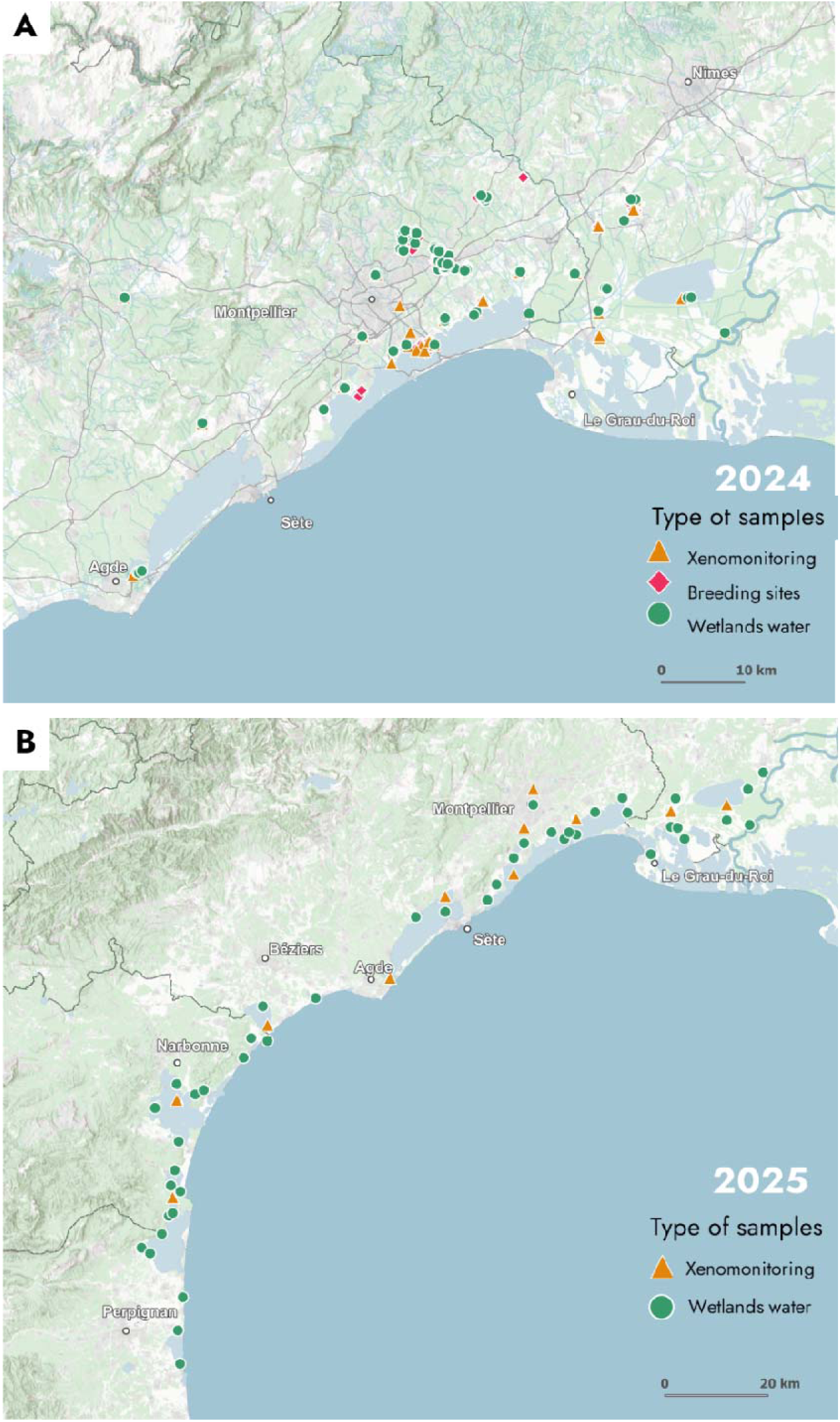
Location map of MX and water samples for 2024 (A) and 2025 (B) campaigns in Occitania, France.

In addition to routine surveillance, 27 MX traps were opportunistically deployed in 9 municipalities during vector control operations in response to confirmed local arbovirus cases. These reactive samplings followed the same standardized protocol, using the same baiting system and adapters, with traps usually active for 3 to 4 days.

Collections in repeatedly sampled wetlands were conducted between the 29^th^ of April and the 8^th^ of October 2024, at 22 spots, and 14 additional wetlands in the vicinity of arbovirose cases were sampled during vector control operations. Waters from artificial breeding sites were collected between the 5^th^ of July and the 14^th^ of October 2024, in 10 different municipalities plagued by confirmed local arboviral circulation.

#### 3.2.2 2025 practical application to regional monitoring

A sampling network of 45 wetlands and 11 mosquito traps equipped with MX adapters was deployed across Occitania, covering ca 200km of the Mediterranean coastline, at fixed locations throughout the sampling season (Fig.2B), from the 1^st^ of April to the 23^rd^ of October 2025.

The wetlands, distributed across 29 municipalities, presented various hydrologic regimes (permanent, temporary or regulated), management intensities (from natural systems to heavily managed sites such as salt marshes), salinity level (freshwater, brackish or saline), but all followed the same criteria as for the 2024 campaign: avian biodiversity hotspots, nesting and migratory stopover zones for wild birds, favourable habitats for Culex mosquitoes.

The 11 mosquito traps with MX adapters were systematically powered by mains electricity to enable continuous functioning. For sites open to the public, the traps were set in metal enclosure BG-Urban Trap stations (Biogents, Germany). Since Culex was specifically targeted, the attractant system was set up accordingly: the CO₂ was released from 10L pressurized cylinders between 6:45 and 8:00 and then, between 19:00 and 00:30 (Veronesi et al., 2012).

MX adapters and water samples were collected biweekly, providing up to 14 samples over the 28-week study period. To best reflect the reactive and operational nature of this surveillance methodology, all field sampling, trap collection, transportation, molecular analyses, and data interpretation were completed within one week.

### 3.3 Environmental water and MX analyses

#### 3.3.1 Development and test of West-Nile and Usutu viruses dPCR detection system and specificity validation on target and non-target viruses

The dPCR detection system for WNV was adapted from Kolodziejek et al. (Kolodziejek et al., 2014), targeting the 5’-UTR region. The probe was used as described, but primers were re-designed to be more specific to WNV. The USUV detection system was adapted from Jöst et al. (Jöst et al., 2011), targeting the NS1 region. As for WNV, primers were re-designed to be more specific to USUV. For both viruses, sequences analysis and oligonucleotides designs were conducted using the Muscle algorithm for alignments (Edgar, 2004) and Primer3 primer design program on Primer3plus website (Untergasser et al., 2007). In silico specificity of primers and probes was evaluated using nucleotide BLAST (Camacho et al., 2009) against GenBank database (Sayers et al., 2024).

To assess the ability of primers and probes to specifically target WNV and USUV, RT-dPCR amplification was conducted on RNA extract from both viruses. Non-target viruses were also chosen for specificity assays : (i) WNV and USUV to test cross-reactivity between both targets, (ii) other Arboviruses as dengue (DENV-1, DENV-2, DENV- and DENV-4), Chikungunya and Zika viruses, (iii) viruses commonly monitored in water-based surveillance as Hepatitis A virus (HAV), Measles virus (Wild-type strain), Respiratory syncytial virus A and B (RSV-A and B), SARS-CoV-2, Influenza A virus, H5N1 avian influenza virus and Norovirus GI/GII.

#### 3.3.2 Digital-PCR method characterization and quality control set up

A multiplex assay was developed to quantify WNV and USUV in the same analysis. Two other targets were incorporated in the multiplex as controls. First, a multispecies mosquito probe (hereafter called “pan-mosquito”; designed from a consensus sequence of 146 ITS1 sequences from genera Aedes, Culex and Anopheles). Then, a system targeting a random exogenous RNA that was added in small amount to each sample in order to determine the presence of inhibitors (hereafter called “inhibition control”). Positive and negative controls of dPCR were also added to analysis. Digital-PCR reactions were conducted using a multiplex assay on a QIAcuity® Eight Plateform System (Qiagen, Germany), using the QIAcuity ® One-Step Viral RT-PCR Kit (Cat. No. 210212) and QIAcuity Nanoplate 26 K 24-wells (Cat. No. 250001). The dPCR reaction mixtures were prepared in a preplate as follows. For each reaction, 10 µL of 4 × One-Step Viral RT-PCR Master Mix, 0.4 μL of 100 × Multiplex Reverse Transcription Mix, 1 μL of the primers/probes mix (final concentration 0.45 µM of both primers and 0.125 µM of probe), and a volume of sample nucleic acids extract, were combined with H_2_O up to 40 µL final reaction volume. Reaction mixtures were then transferred into a QIAcuity® Nanoplate and the plate was loaded onto the fully automated QIAcuity ® Eight instrument. The workflow included (i) priming and rolling step to generate and isolate the chamber partitions (26,000 partitions), (ii) amplification step under the following cycling protocol: 50 °C for 40 min for reverse transcription, 95 °C for 2 min for enzyme activation, 95 °C for 5 s for denaturation, and 58 °C for 60 s for annealing/extension for 40 cycles; and (iii) imaging step by reading fluorescence emission after excitation of the probe at the appropriate wavelength.

Samples from the 2024-campaign were first analysed using 2 µL of sample both for MX samples and water samples. This input volume was applied to all samples collected from the beginning, up to those sampled in mid-September. An optimisation of the input volumes was then performed on chosen samples (varying in term of origin) to test potential PCR inhibition and improve the representativity of the measure in the remaining samples from 2024 and 2025 campaign. For MX samples, USUV-spiked samples at different concentrations (Low: 4.6 TCID_50_.mL^-1^, Medium: 4.6 x 10^1^ TCID .mL^-1^ and High : 4.6 x 10^2^ TCID .mL^-1^) were prepared, extracted and tested in duplicates with three different input volumes (2, 4 and 8 µL). For water samples, artificial breeding sites were used, as they exhibited the highest levels of inhibition in preliminary tests. Two water samples (Zoological park of Montpellier and artificial breeding sites from vector control operation in Pérols, south Montpellier) were tested in triplicates using two sample input volumes (4 and 6 µL). Pan-mosquito and inhibition controls concentrations were compared depending on the sample input volume.

The smallest copy number of the target sequence detectable in a reliable manner per dPCR well (absolute limit of detection; LoD) was determined for WNV and USUV. A positive internal control was diluted to achieve final concentrations corresponding to, respectively, 5 and 10 RNA copies per dPCR reaction (copies deposited in plate wells). Each dilution was measured in ten replicates on the same dPCR run. Moreover, the smallest copy number of the target sequence from which the abundance of that sequence can be measured accurately and reliably (absolute limit of quantification; LoQ) was measured for WNV and USUV by analysing a series of positive control dilutions and identifying the lowest concentration at which quantification remained accurate and reproductible, based on inter-replicate variance and established method acceptance criteria. For this purpose, ten independents’ solutions were prepared from each positive control (WNV or USUV) to reach a final concentration of 33.33 RNA copies/µL of reaction mix for each target, and each dilution was analysed once, resulting in ten dPCR measurements per tested concentration level, performed on four different thermocyclers. The blank limits (LoB), i.e. the highest concentration measured in the absence of the target in the dPCR well, were determined. To characterize this “background noise” of the method, thereby ensuring no false positive could be detected, two extraction negative controls which followed the entire process of sample acid nucleic extraction were quantified in ten replicates on two different thermocyclers. For each target of the multiplex, the median was calculated from the obtained twenty concentrations (in copy.µL of dPCR reaction^-1^) and standard deviation was determined. The LoB was then calculated for each target using the formula:

LoB = median + 1.645 × sd

With sd the standard deviation, and the 1.645 factor corresponding to the 95th percentile of a normal distribution. This threshold indicates that 95% of measurements obtained from negative controls are expected to fall below the LoB, and that only 5% of values would exceed the LoB due to random variation, corresponding to a low probability of false-positive detection.

#### 3.3.3 Method development for acid nucleic extraction from MX samples

A preliminary approach –hereafter called the ‘alternative’ protocol– was first implemented for the processing of MX samples derived from mosquito excreta using an inhouse-protocol developed by IAGE. Mosquito excreta were manually collected using a loop and resuspended in 0.2 mL of 1X Phosphate-Buffered Saline pH 8 by shaking. The entire volume of solubilized excreta was then directly used for DNA/RNA extraction performed with the Indimag Pathogen Kit (Indical Bioscience, Germany).

In parallel, an optimized protocol was developed, adapted from the method described by Bigeard et al. (Bigeard et al., 2024) –hereafter called the ‘literature’ protocol–, to improve mosquito excreta recovery. Briefly, the aluminium foil from the MX adapter was coiled and placed in a 50 mL plastic tubes (Greiner Bio-one, Cat. No.227261). Sample was then soaked in a 1.2 mL VXL lysis buffer (Cat. No.1069974, Qiagen, Germany) and vortexed for 5 min. The supernatant was recovered, and 0.2 mL were used for downstream DNA/RNA extraction using the Indimag Pathogen Kit (Cat. No. SP947457, Indical bioscience, Germany). Nucleic acid were then eluted in 0.1 mL H_2_O.

Efficiency of ‘literature’ and ‘alternative’ pre-treatments for viral recovery and detection sensitivity was compared by side-by-side analyses. To do so, mosquito excreta were artificially deposited onto aluminium foil substrates and spiked with USUV at three different concentrations: Low (4.6 × 10² TCID₅₀.mL^-1^), Medium (4.6 × 10³ TCID₅₀.mL^-1^) and High (4.6 × 10⁴ TCID₅₀.mL^-1^). Each concentration level was prepared in duplicate to allow independent nucleic acid extraction by each institute. Viral detection and quantification were performed using dPCR. The resulting data were subsequently compared across protocols. Based on these comparative results, the ‘literature’ protocol was selected for further analyses of the remaining mosquito excreta samples from 2024 and samples from the 2025 field application.

#### 3.3.4 Method development for acid nucleic extraction from wetlands and artificial site water samples

The water samples, with volumes comprised between 0.1 to 1 L, were kept at 4°C after sampling until nucleic acid extraction (Fig.1C). Extractions of total DNA/RNA for dPCR analyses were performed following the IAGE patented method (Durandet et al., 2022) with an optimized pre-treatment. Briefly, the patented method consists in thoroughly mixing the total volume of water, sub-sampling and applying 15 mL from each homogenized sample to an Amicon® Ultra-15 Centrifugal Filter Unit (cut-off: 10 kDa, Cat. No. UFC901096, Merck Millipore, Germany), previously hydrated with 2.5 mL of MQ water and spin for 10 min at 3234 g, at 4°C. The amicon was then centrifuged for 35 minutes at 3234 g, at 4°C. If the resulting volume was inferior to 0.5 mL, the tube was centrifuged again for 10 minutes at 3234 g, at 4°C.

The optimized version added a centrifugation of the 15 mL sub-sampling for 15 min at 3234 g at 4°C. For each sample, pellet was conserved on ice until further treatment. Supernatant was recovered and applied to an Amicon® Ultra-15 Centrifugal Filter Unit (cut-off: 10 kDa, Cat. No. UFC901096, Merck Millipore, Germany) as for the patented method. The resulting volume of the concentration was used to resuspend the corresponding pellet for each sample. Then, 0.4 mL of Nucleo Protect VET Buffer (Cat. No. 740750, Macherey Nagel, Germany) was added before grinding with two 3-mm stainless steel beads on a Geno/Grinder ® instrument (Spex Sample Prep LLC, USA) for 15 s at 1500 cpm, thrice. In both protocols, extractions were then performed from 0.2 mL using the Indimag Pathogen kit (Cat. No. SP947457, Indical Bioscience, Germany) and nucleic acid were then eluted in 0.1 mL H_2_O.

To compare the patented method with the optimized version, two water samples and two larval habitat water samples were spiked with USUV at a concentration of 1.10⁷ copies·L⁻¹. Each sample underwent both extraction protocols. Additionally, one non-spiked sample from the Zoological park of Montpellier was extracted in triplicates using both methods; only pan-mosquito concentration, was quantified and used as a positive control for comparison. All samples, from 2024-proof of concept and 2025-field trial, were subsequently analysed with the optimized version.

Because such sampling campaigns may require reanalysis of previously analysed samples, and given the complexity of water as a matrix, we evaluated the stability of viral targets during raw water storage. Thus, we assessed whether storage at 4°C reliably preserve viral RNA and determined the maximum duration over which this stability could be ensured. Two WNV-positive samples–one from wetlands and one artificial site water– were stored at 4°C and re-extracted at 8, 15, 23, 25 and 28 days post-collection.

### 3.4 Virus sequencing

Whole-genome sequencing of the virus was performed for all MX or mosquito samples with a sufficiently high viral load (Ct < 32). PCR amplifications were conducted using eight pairs of overlapping primers targeting segments of the region of interest (Bigeard et al., 2024). Reverse transcription and amplification were carried out using the SuperScript IV One-Step RT-PCR kit (ref. 12594100, Thermo Fisher, USA). Each 25 μL reaction contained 12.5 μL of 2X Master Mix, 1.25 μL (10 μM) of each forward and reverse primer, 6.5 μL of RNase-free water, 0.5 μL of RT enzyme mix, and 3 μL of extracted genetic material. Thermal cycling conditions were as follows: 55°C for 10 min, 98°C for 2 min, followed by 40 cycles of 98°C for 10 s, 55°C for 10 s, and 68°C for 1 min 45 s, with a final extension at 68°C for 5 min. PCR products were visualized by electrophoresis on a 1% agarose gel stained with EZ-Vision DNA dye (ref. 1B1680, VWR Chemicals, USA) to verify amplification success and amplicon size.

Amplicons were then purified and sent to the Unité des Virus Émergents (UVE, Aix-Marseille University, France) for downstream processing. Equimolar quantities of each amplicon were pooled and quantified using a Qubit fluorometer (Thermo Fisher, USA). DNA fragmentation into ∼250 bp fragments was performed with a Bioruptor sonicator (Diagenode, Belgium). Library preparation was carried out using the Ion Plus Fragment Library Kit (ref. 4471252, Thermo Fisher, USA) and the automated AB Library Builder system (ref. 4463794, Thermo Fisher, USA). The pooled libraries were quantified by qPCR to confirm equimolarity, then amplified by emulsion PCR and loaded onto an Ion Chip using the Ion Chef instrument (Thermo Fisher, USA). Sequencing was performed on the Ion Torrent S5 system (Thermo Fisher, USA), generating high-quality reads for downstream analysis. Following sequencing, raw reads were demultiplexed based on their barcode identifiers, quality-filtered, and aligned to the closest viral reference genome. Consensus sequences were assembled from well-covered regions, while poorly covered areas were masked with “N” to denote undetermined nucleotides.

In parallel to the sequencing performed for the identification of circulating lineages, Sanger sequencing was conducted on samples with low WNV signals in dPCR to confirm that the detected target corresponded to the virus. Total nucleic acid from selected samples were subjected to DNase treatment using TURBO™ DNase (ref. 10722687, Thermo Fisher, USA) to remove residual genomic DNA prior to downstream analyses. For each reaction, a 10 µL mix was prepared containing 1 µL of 10X TURBO™ DNase Buffer, 1 µL of TURBO™ DNase, 1 µL nuclease free-water and 8 µL of nucleic acid extract. Samples were mixed and incubated for 30 min at 37 °C. DNase inactivation was performed by adding 3.6 µL of 50 mM EDTA (final 10X dilution of the supplied 0.5 M EDTA) followed by incubation at 75 °C for 10 min. RNA was then reverse transcribed. For each sample, a reaction volume of 20 µL was prepared, consisting of 4 µL LunaScript® RT SuperMix (ref. E3010L, New England Biolabs, USA), 4 µL nuclease-free water, and 12 µL of DNase-treated extract. Reverse transcription was performed under the following cycling conditions: 25 °C for 2 min, 55 °C for 10 min, and 95 °C for 1 min. cDNA concentration was quantified using sDNA High Sensitivity or Broad Range assays (Thermo Fisher, USA). Targeted amplification of virus was performed using the Q5® Hot Start High-Fidelity 2X Master Mix (ref. M0494S, New England Biolabs, USA) with the specific forward and reverse primers. PCR programs were adapted according to primer annealing temperatures previously optimized by dPCR. Amplicons were checked by agarose gel electrophoresis (2% TBE), visualized under UV illumination, and purified using the QIAquick PCR Purification Kit (QIAGEN, Germany). Purified amplicons were quantified by Qubit™ dsDNA BR assay, diluted to the required concentration, and prepared for Sanger sequencing using the Eurofins Mix2Seq service. For each sample, 5 µL of purified DNA (1 ng.µL^-1^) and 5 µL of either forward or reverse primer (5 pmol.µL^-1^) were submitted following provider specifications.

### 3.5 Data processing

Raw data generated by the QIAcuity ® Software suite V3.1.0.0. platform were interpreted based on the performance of internal and external dPCR controls. Positive dPCR controls for WNV and USUV were included at both high and low concentrations (3xLoD), to verify assay sensitivity, while the inhibition control was added to all samples and run alone at 3xLoD. Negativity of extraction and analytical control was checked. When acceptance criteria were met and at least 20,000 valid partitions were obtained, we set fluorescence threshold using positive controls and transformed raw partition counts into numbers of copies per mL, adjusting for the acid nucleic extraction process. Samples showing inhibition were re-analysed with a 1:10 dilution. Resulting concentrations were used for downstream statistical and spatial analysis (Fig.2D).

Environmental and mapping data were collected in the field using the “MerginMaps” mobile application v2025.8.0 (Cavallini et al., 2022) on smartphone and processed with QGIS v 3.34.10-Prizren (Nyall Dawson et al., 2025). All quantitative results were integrated into a centralized database, enabling structured data exploration. All analyses were carried out in R v. 3.6.1 (R Development Core Team, 2019) using the packages reshape2 (Wickham, 2007), dplyr (Wickham et al., 2019), lubridate (Grolemund & Wickham, 2011), data.table (Barrett et al., 2025) and rnaturalearth (Massicotte et al., 2025) for the analyses and data manipulation, and ggplot2 (Wickham, 2016) for data visualisation. OpenAI’s ChatGPT (version GPT-4.0, accessed January 2026) was used for R code improvement.

## 4. Results

### 4.1 Development and characterization of a digital-PCR method targeting WNV and USUV

#### 4.1.1 Development of a reliable multiplex assay

The dPCR assays for WNV and USUV, consisting in two primers and a TaqMan probe, were initially validated in silico through sequence alignment and BLAST analysis to assess theoretical specificity. No significant matches with non-target viruses were detected. Designed oligonucleotids (Supplementary Tab. S2) amplified a fragment of 105 bp for WNV in the 5’UTR region and a fragment of 88 bp for USUV in the NS1 region (Supplementary Fig. S1). Each detection system specifically amplified its respective positive control in simplex assays without generating any signal for non-target viruses. No cross-reactivity was observed neither between WNV and USUV nor with any other tested viruses. The multiplex assay, comprising WNV, USUV, Pan-mosquito and the inhibition control, was considered validated as the dPCR quantification of each target in multiplex did not decrease by more than 10% compared to the corresponding simplex assay, confirming that simultaneous detection did not compromise assay sensitivity or accuracy.

Inhibition tests were conducted to enhance the representativity of the measure by increasing sample volume input in the dPCR reaction mix, while not inducing a loss in quantification. For MX samples, the comparison was performed on USUV-spiked samples. A detectable signal was observed for all USUV concentrations and volumes tested except for the 2 µL input which did not allow detection of the lowest USUV spiking concentration (Fig. 3a). This indicated that low viral load could be reliably detected preferably with 4 or 8 µL inputs. An input volume of 8 µL was selected as no quantification decrease was observed for the three spiking concentrations tested. For water samples, two artificial breeding sites were chosen, and pan-mosquito and inhibition control concentrations were compared (Figure 3b). Increasing the dPCR input volume did not affect the inhibition control detection but resulted in a lower final quantification of the pan-mosquito target for both water samples. Therefore, an input volume of 4 µL was selected for water samples.

**Figure 3:**
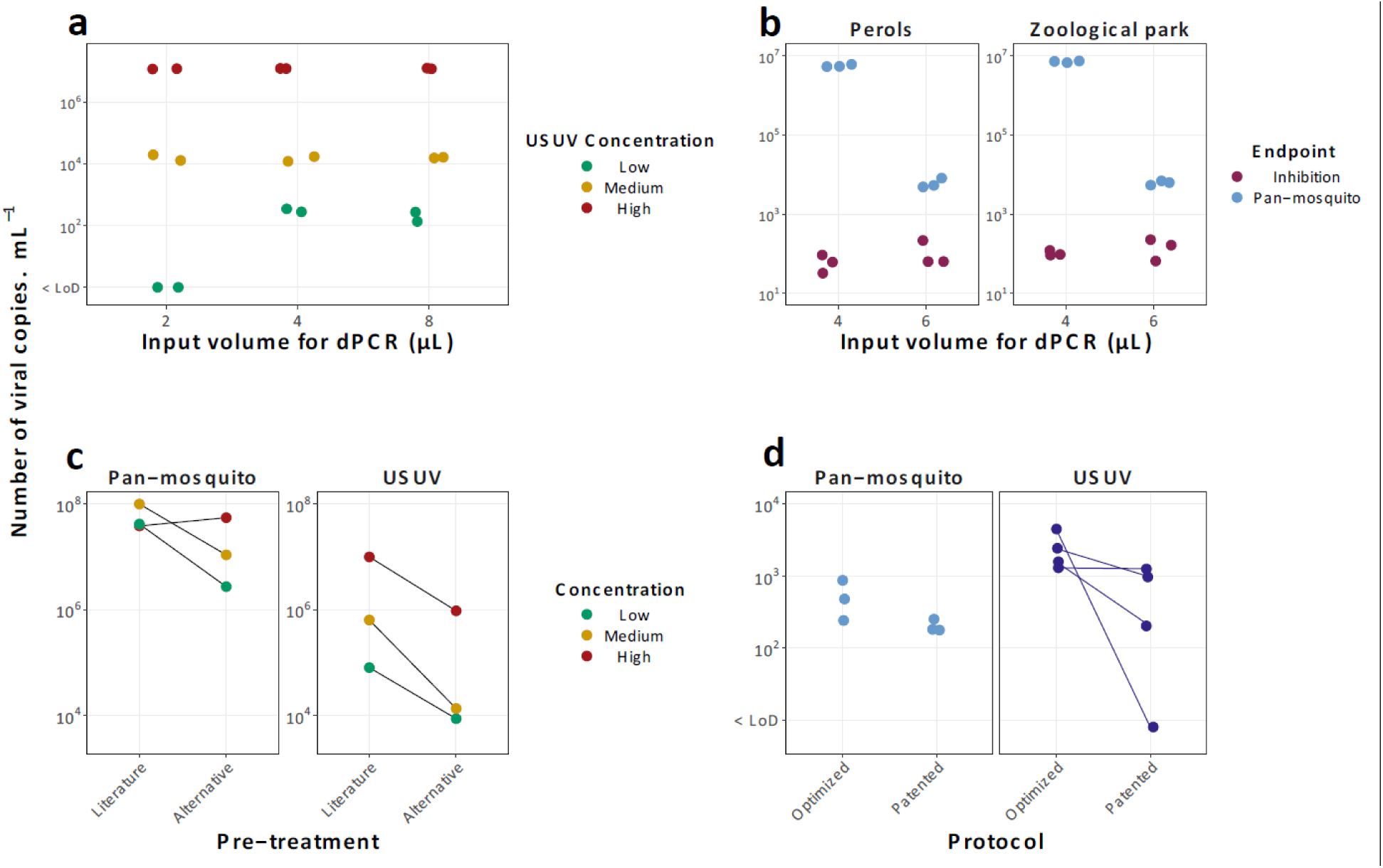
Digital-PCR and acid nucleic extraction methods optimization from mosquito excreta and water samples. dPCR inhibition assays were conducted both from (a) USUV-spiked MX samples, with 3 different sample input volumes (Mosquito excreta samples were spiked at Low : 4.6 TCID₅₀·mL⁻¹, Medium : 4.6 × 10¹ TCID₅₀·mL⁻¹, and High : 4.6 × 10 TCID₅₀·mL⁻¹ concentrations), and (b) 2024 campaign artificial breeding site samples (Pérols and Zoological Park), with 2 sample input volumes tested, for water analysis. Water samples were not spiked, only pan-mosquito and inhibition control concentrations are shown. Extraction methods optimizations were conducted in (c) a side-by-side analysis (‘literature’ protocol described in Bigeard et al. (2024), vs the alternative one) on three USUV-spiked MX samples at 3 different concentrations (Low : 4.6× 10^2^ TCID .mL⁻¹, Medium : 4.6× 10^3^ TCID .mL⁻¹, and High: 4.6× 10^4^ TCID .mL⁻¹), and for water samples, on (d) four USUV-spiked samples at a concentration of 1.10⁷ copies·L⁻¹ processed either with the patented or the optimized protocol, in addition to a non-spiked sample analysed in triplicates using both methods (only results for pan-mosquito target are shown). For all graphs, Y-axis is in logarithmic scale log10(x+1). “< LoD” = below the limit of detection.

#### 4.1.2 Analytical Performance

The analytical performance of the multiplex dPCR assay was measured by determining the LoB, absolute LoD, and absolute LoQ for WNV, USUV, and the pan-mosquito target (Tab. 1). LoB values were low for all detection systems, reflecting minimal background signal and allowing a clear distinction between true positive reactions and false-positive events. The LoB was slightly higher for the pan-mosquito target than for WNV and USUV but remained within an acceptable range for rare-target detection. The absolute LoD for both WNV and USUV was 0.125 copies per reaction, corresponding to the lowest concentration at which at least 90% of dPCR reactions yielded a positive signal. This demonstrates the high analytical sensitivity of the assay and its suitability for detecting low viral loads in environmental and MX samples. The absolute LoQ for WNV, USUV, and the pan-mosquito target was determined to be 1.25 copies per reaction. At this concentration, quantitative results met the predefined criteria for repeatability (RSDr ≤ 25%) and accuracy (±30% of the expected theoretical copy number). Compliance with the dMIQE 2020 guidelines(dMIQE Group & Huggett, 2020) was ensured and the corresponding checklist is provided (Supplementary Tab. S3).

**Table 1.** Analytical performance of the WNV - USUV – Pan-mosquito multiplex assay. Limits are expressed in number of viral copies of target per µL dPCR reaction. LoD : limit of detection, LoQ : limit of quantification, LoB : limit of blank.

| Target | Absolute LoD | Absolute LoQ | LoB |
| --- | --- | --- | --- |
| WNV | 0.125 | 1.250 | 0.021 |
| USUV | 0.125 | 1.250 | 0.021 |
| Pan-mosquito | 0.125 | 1.250 | 0.030 |

### 4.2 Development of a reliable process for the analysis of environmental samples

#### 4.2.1 Comparison of MX nucleic acid extraction protocols

Comparative tests were conducted to evaluate the performance of two nucleic acid extraction protocols for MX samples: the protocol described in the literature(Bigeard et al., 2024) and an alternative in-house protocol. Each participating institution processed the samples using its respective pre-treatment and extraction workflow, either applying the protocol described in the literature or the alternative in-house protocol. The pan-mosquito target was used as an extraction control to assess nucleic acid recovery efficiency, while USUV quantification reflected viral detection performance. For comparable spiking levels, both pan-mosquito and USUV signals were consistently higher when using the protocol described in the literature compared to the alternative method (Fig.3c). This demonstrates a superior extraction and detection efficiency of the literature-based pre-treatment for this type of sample. Based on these results, the protocol described in the literature was selected for further analyses. To improve the solubilisation of mosquito excreta, a minor adaptation was implemented by increasing the lysis buffer volume from 0.8 mL to 1.2 mL. This modified version was validated and subsequently retained for routine processing of MX samples.

#### 4.2.2 Comparison of water nucleic acid extraction protocols

A comparative evaluation was performed between the in-house protocol extraction protocol, originally developed for SARS-CoV-2 detection in wastewater, and an optimized protocol adapted for the detection of arboviral targets in wetland and artificial breeding site water matrices. The optimized protocol resulted in higher detection levels for both USUV in spiked-wastewater samples and pan-mosquito target in the larval habitat sample compared to the patented method but one (Fig.3d). These results indicate an improved recovery and detection efficiency of viral and mosquito-derived nucleic acids using the optimized extraction workflow for environmental water matrices. The stability test performed on the water samples showed no decrease of WNV and pan-mosquito quantification after nucleic acid extraction up to 15 days post-collection (Supplementary Tab. S4), ensuring reliable storage for subsequent analysis. Consequently, the optimized protocol was selected and applied to all water samples analysed in the 2024 proof-of-concept and the 2025 field campaigns.

### 4.3 WNV lineage identification classification and confirmation of dPCR assay specificity by sequencing

Over the two-year sampling period, we obtained seven specimens with WNV viral loads high enough to enable whole-genome sequencing (Tab.2). Two genomes originated from MX samples, five from individual mosquitoes collected in traps, and four from water samples. So far, all sequences analysed belonged to lineage 2. In 2024, WNV-infected mosquitoes were detected even though the MX samples were negative. This discrepancy was no longer observed in 2025, likely due to the optimized MX screening protocol implemented that year.

**Table 2:** Metadata for WNV sequences obtained from high positive samples in 2024 and 2025. Matrices were excreta collected by xenomonitoring, water from wetland or artificial breeding site, or entire trapped mosquito individuals. *ND stands* for “Not Determined”, *NA* for “Not applicable”.

| Accession<br>number | Sampling<br>date | Matrix | Site type | Municipality | Lineage | Sequencing<br>method | Genome<br>coverage<br>(%) | Sequence<br>identity<br>(%) |
| --- | --- | --- | --- | --- | --- | --- | --- | --- |
| OZ263592 | 30/09/2024 | Mosquito excreta | Natural reserve | Mireval | 2 | WGS | 51,11 | NA |
| OZ263593 | 17/09/2024 | Mosquito | Zoological park | Montpellier | 2 | WGS | 84,64 | NA |
| OZ263594 | 13/09/2024 | Mosquito | Marsh | Mauguio | 2 | WGS | 93,6 | NA |
| OZ263595 | 16/08/2024 | Mosquito | Marsh | Saint-Laurent-d'Aigouze | 2 | WGS | 93,6 | NA |
| OZ263596 | 30/08/2024 | Mosquito | Marsh | Saint-Laurent-d'Aigouze | 2 | WGS | 93,6 | NA |
| Ongoing | 29/07/2025 | Mosquito excreta | Zoological park | Montpellier | 2 | WGS | ongoing | NA |
| Ongoing | 09/09/2025 | Mosquito | Educational farm | Saint-Jean-de-Védas | 2 | WGS | ongoing | NA |
| NA | 09/09/2024 | Water (Artificial site) | Residential area | Baillargues | ND | Sanger | NA | 98,4 |
| NA | 09/04/2025 | Water (Wetland) | Saltmarsh | La Palme | ND | Sanger | NA | 98,5 |
| NA | 09/04/2025 | Water (Wetland) | Marsh | Narbonne | ND | Sanger | NA | 100 |
| NA | 09/04/2025 | Water (Wetland) | Temporary brackish lake | Fleury | ND | Sanger | NA | 96,9 |

Sanger sequencing was performed on selected WNV-positive samples to confirm the specificity of the dPCR assay and that the detected signal corresponded to WNV. For two MX samples from 2024 showing high WNV concentrations in dPCR (≈80 copies·µL⁻¹ reaction), targeted amplification consistently yielded amplicons of the expected size. The resulting consensus sequences showed unambiguous alignment to West Nile virus reference genomes, with no match to sequences from other viruses. All negative controls remained free of amplification, confirming the absence of contamination. For four low-positive samples (one artificial breeding site from 2024 and three wetlands from 2025; ≤1 copy·µL⁻¹ reaction), sequencing also produced high-quality reads aligning to WNV. Although the amplified region is known to share partial homology with USUV, the most divergent segment between the two genomes (positions 20–50 bp and 60–100 bp within the amplicon) was fully resolved. In both regions, nucleotide identity was consistent with WNV and not with USUV, ruling out a cross-amplification. Additional verification confirmed the absence of homology between the amplicon and the Culex pipiens genome, in agreement with the lack of reported viral integrations in this species. Altogether, these results demonstrate that the dPCR assay specifically detects WNV RNA, even at very low target concentrations, and does not produce nonspecific amplification from related viruses or host genomic material.

### 4.4 2024 operational proof of concept

Overall, WNV was detected 29 time in mosquito excreta and waters, in 11 different municipalities. From the repeatedly sampled sites, 9 out of 139 MX traps and in 8 out of 171 wetland water samples were positive, providing respectively up to 2.2.10 and 9.8.10 copies per mL (Figs.4 and 5). It was also detected during vector control operations in 1 out of 15 water samples from surrounding wetlands, 1 out of 32 MX traps and in 10 out of 45 artificial breeding sites.

The earliest WNV detection happened on the 1 of July 2024 in a mixture of water from wetlands and poultry ponds in the educational farm of Saint-Jean-de-Vedas. From the repeatedly sampled sites, WNV was last detected the 30 of September at the same location, and in wetlands samples from the Natural Reserve in Mireval. One MX sample was also WNV positive the 1 of October, during a vector control operation around the city of Vauvert. Comparatively, the first detection in human blood was the 9 of August(Grard et al., 2025), the first alerts of WNV autochthonous human case, the 3 and the 24 of September for Hérault and Gard(Ministry of Health, Family, Autonomy and Disabled Persons, 2026), and the first equine alerts of WNV, the 6 and the 13 of September(RESPE, 2024a, 2024b).

USUV was detected nine time in six municipalities. It was present in one MX trap –3.2.10 copies per mL– and in six wetland water samples –up to 62.5 copies per mL– belonging to the repeatedly sampled sites, with a first detection the 9th of July at the Zoological park of Montpellier (Figs. 4 and 5). It was also detected during vector control operations in two artificial breeding sites, including the earliest detection the 5 of July in Pérols.

**Figure 4:**
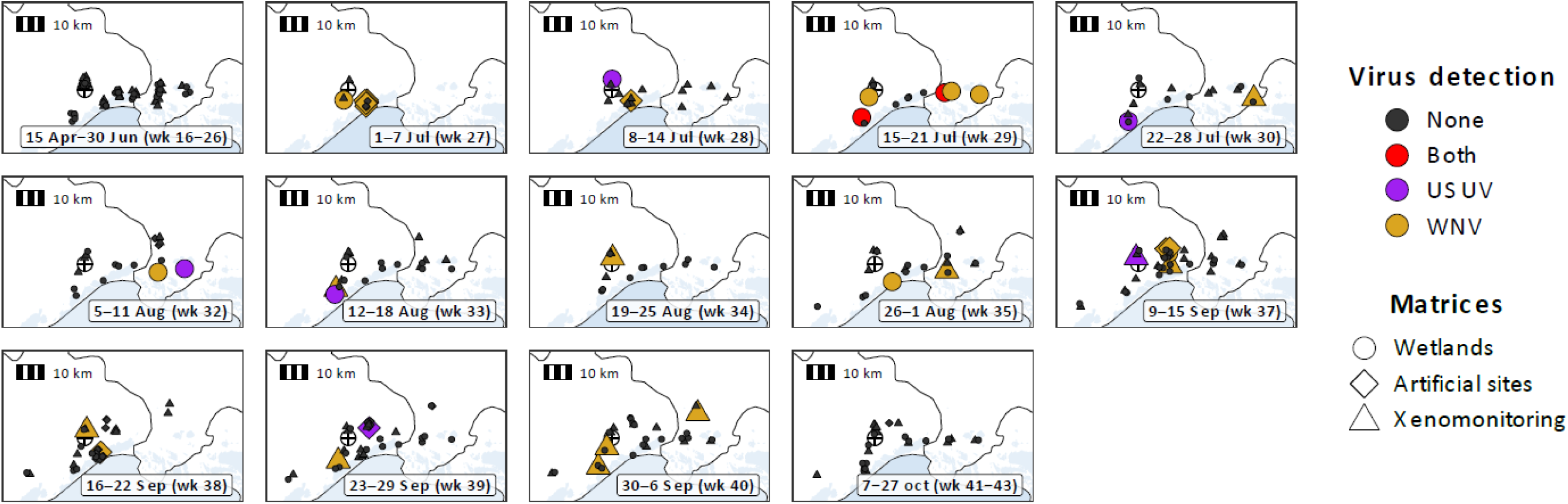
maps of viral detection over the sampling season 2024 in wetland waters, mosquito excreta in MX adapters through xenomonitoring, or artificial breeding sites. The circled plus (⊕) symbolises the city of Montpellier. Overlapping points slightly jitter for graphical purposes.

**Figure 5:**
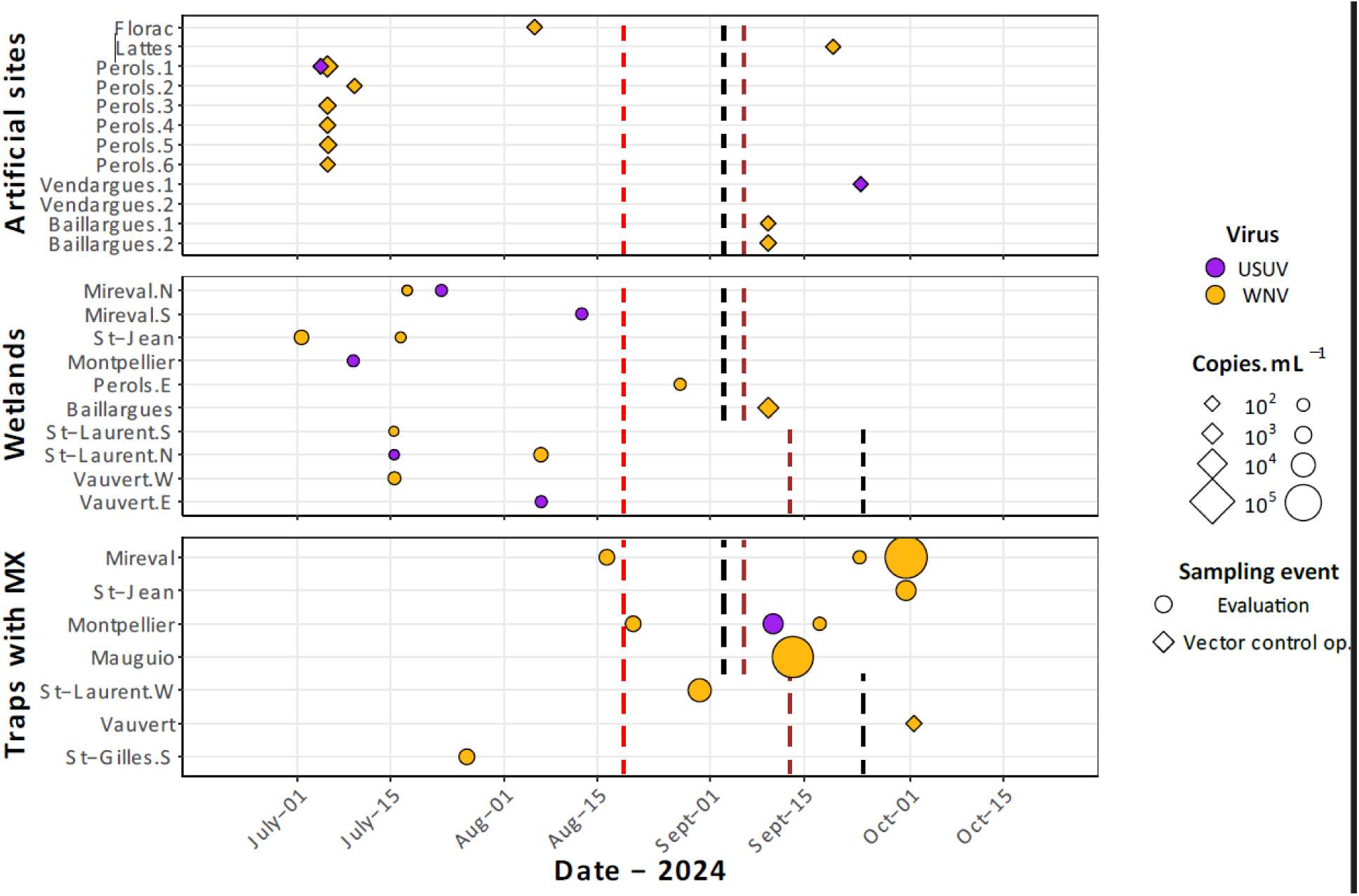
Number of viral copies of WNV and USUV per mL in artificial breeding sites (upper panel), wetland waters (middle panel) and in mosquito excreta collected in traps equipped with MX adapters (lower panel), from West (Mireval/Florac, top) to East (Baillargues, Vauvert and St-Gilles, bottom), regularly sampled from evaluation campaign or opportunistically collected during vector control operations, over the 2024 sampling season. The dashed lines indicate the first human detection of WNV in blood donations in Occitania (in red), the first alert of human autochthonous case of WNV (in black) and the first equine alert of WNV (in brown) in their respective French departments in 2024. Only sites with at least one positivity of either of the viruses are shown.

WNV and USUV were simultaneously detected in one artificial breeding site during vector control operation in Pérols the 5^th^ of July, in the Petite Camargue wetlands (Vauvert municipality) the 15 of July, and in the wetlands of the Natural Reserve of Mireval the 17 of July.

### 4.5 2025 real-life campaign of regional monitoring

WNV was detected in 76 water samples over the 28-week study period, collected from 40 distinct sites between the 9^th^ of April and the 23 of October (Figs 6 and7). Detection was highest at the beginning of the campaign, with 32 and 26 WNV-positive water samples in April and May, respectively, reaching concentrations of up to 1,984 copies·mL⁻¹—the highest levels observed in this matrix. From June, the number of positive water samples declined, ranging from two to seven detections per month until October.

**Figure 6:**
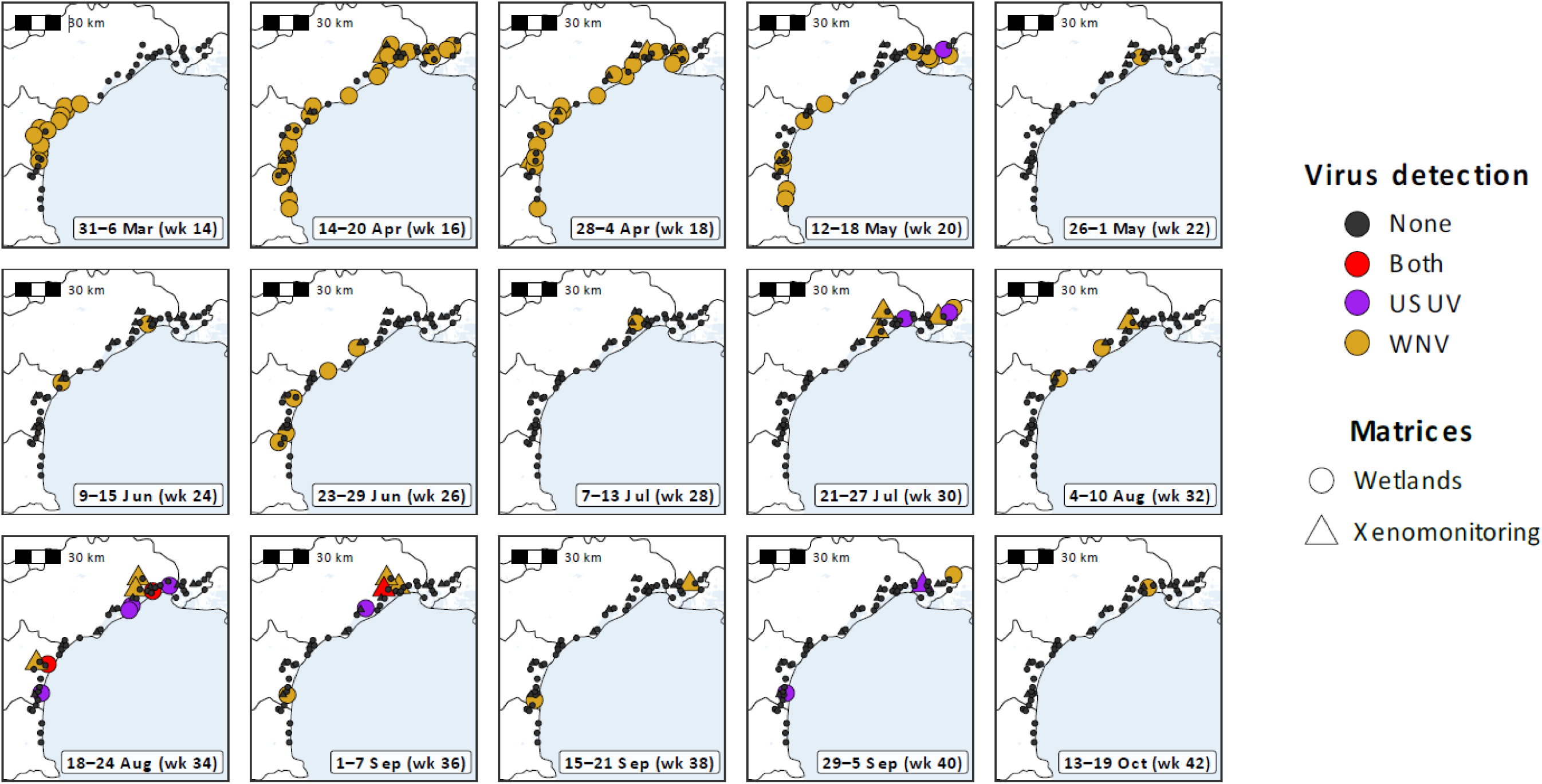
maps of viral detection over the sampling season 2025 in wetland waters or mosquito excreta in MX adapters through xenomonitoring.

Additionally, WNV RNA was detected in mosquito excreta in MX adapters at seven sites, with 14 positive samples identified between the 24^th^ of April to the 23^rd^ of September (Figs 6 and 7). Only one and two samples were positive in April and May respectively, three in July with up to 1.6.10 viral copies per mL, and peaked in August and September with four positive samples each, yielding concentrations up to 3.9.10 and 4.9.10 copies per mL. Interestingly, WNV was detected in several sites already positive in 2024, such as the educational farm of St-Jean-de-Vedas, the Zoological Park of Montpellier, and the Natural Reserves of Mireval and Vauvert. Comparatively, the first detection in human blood in the region was the 25^th^ of August, the only alert of WNV autochthonous human case, the 11 of August in Hérault(Ministry of Health, Family, Autonomy and Disabled Persons, 2026), and the first equine alerts of WNV, the 20^th^ of August in Gard and the 10 of September in Hérault(RESPE, 2024a, 2024b). No such alert occurred in Aude and Pyrénées-Orientales departments.

**Figure 7:**
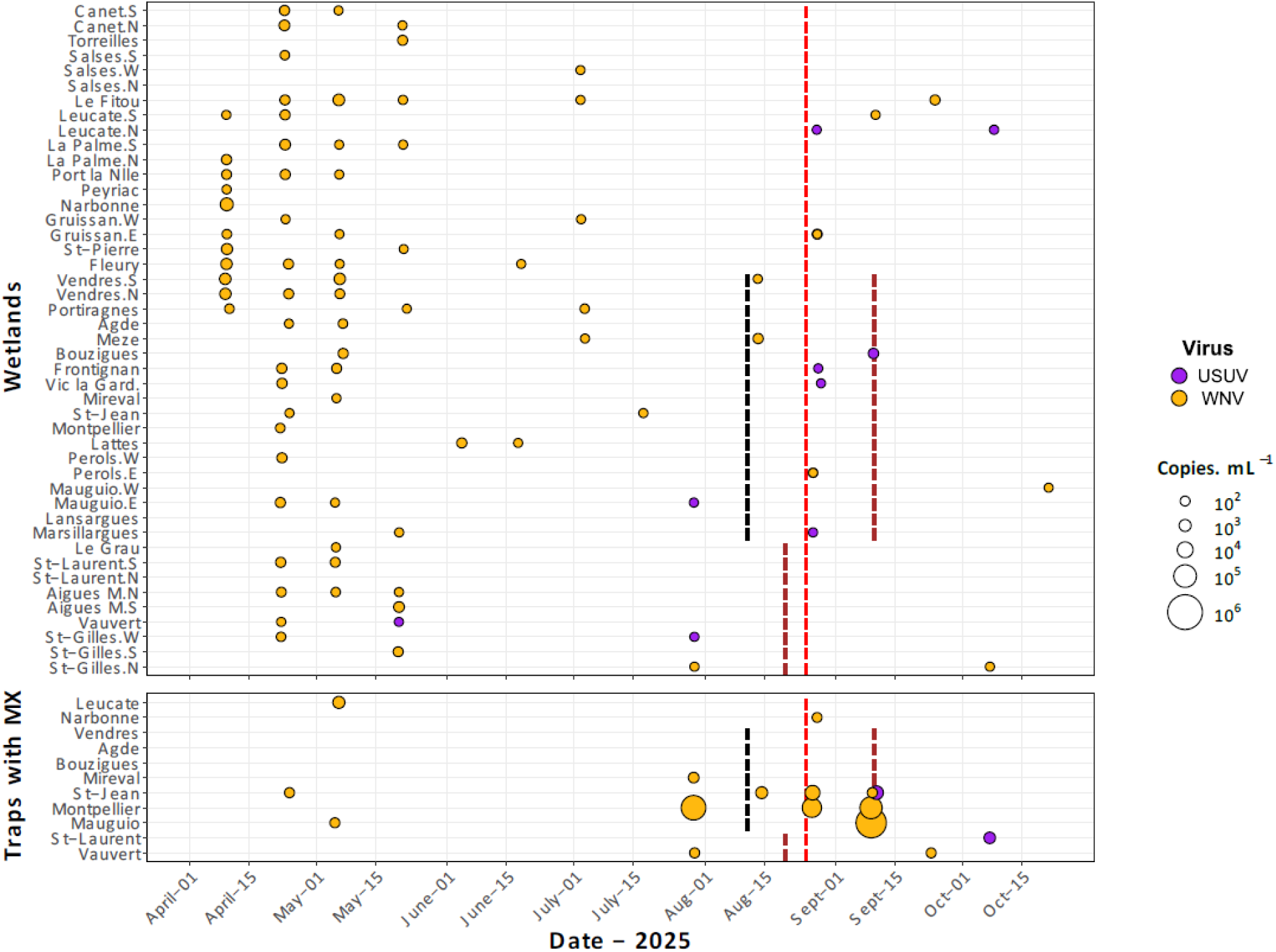
Number of viral copies per mL of WNV and USUV in wetland waters (upper panel), and in mosquito excreta collected in traps equipped with MX adapters (lower panel), from South-West (Canet.S and Leucate, top) to North-East (St-Gilles.N and Vauvert, bottom) over the 2025 sampling season. The dashed lines indicate the first human detection of WNV in blood donations in Occitania (in red), the first alert of human autochthonous case of WNV (in black) and the first equine alerts of WNV (in brown) in their respective French departments (where applicable) in 2025.

USUV was detected 11 times in water samples from 10 sites, between the 20 of May and the 8 of October: once in May, twice in July, peaked in August with 6 positive samples then decreased the following months (Figs 6 and 7). USUV was also found twice in mosquito excreta in late season, the 9 of September in the educational farm, and in the Petite Camargue wetlands of the Natural Reserve of Vauvert (as in 2024) the 7 of October. Up to a 182 copies per mL were found in waters, but excreta yielded a maximum of ca. 3000 copies per mL, in September.

## 5. Discussion

We developed an integrated surveillance strategy combining MX with water sampling from wetlands and mosquito breeding sites, supported by an innovative dPCR assay, enabling early and repeated detection of WNV and USUV during the 2024 and 2025 transmission seasons in Southern France. This multi-matrix approach detected WNV and USUV 29 and 8 times, respectively, among 396 samples in 2024, and 90 and 14 times among 815 samples in 2025, across 22 municipalities. Both years, WNV was identified weeks before autochthonous human cases or equine alerts in the region, highlighting a remarkable temporal precedence decisive for efficient arbovirus surveillance.

The detection of WNV and USUV in environmental waters required substantial methodological developments to ensure analytical specificity, sensitivity, and interpretability of low-level viral signals. A multiplex digital PCR assay targeting the WNV 51-UTR and the USUV NS1 regions was designed to maximize specificity, with no cross-reactivity between WNV and USUV or with a broad panel of non-target viruses. Given the inherent difficulty to interpret low-level detections in environmental matrices, assay performances were characterized using the limit of blank (LoB), the limit of detection (LoD) and the limit of quantification (LoQ), enabling robust discrimination between true positives and background noises. Several RT-qPCR assays optimized for WNV detection report LoD ranging from ca. 1.5 to 15 copies per reaction, depending on the targeted genomic region and viral lineage, demonstrating the high sensitivity achievable with well-designed qPCR assays (Petrović et al., 2007). Multiplex RT-qPCR approaches developed for environmental surveillance, including WNV, report LoDs of approximately 8.8 copies per reaction (Liu et al., 2024) consistent with reliable detection at very low viral loads. For USUV, published RT-qPCR assays report LoDs around 50 genomic copies per reaction in clinical matrices (Cavrini et al., 2011) while optimized surveillance assays have reached detection limits of 11 copies per reaction in mosquito pools (Nikolay et al., 2014). Although these studies do not explicitly define LoB or LoQ, their data on analytical sensitivity provide relevant benchmarks for evaluating assay performance. In comparison, our dPCR system achieved a substantially lower LoD of 0.125 copies per reaction for both WNV and USUV, reflecting the inherent advantage of dPCR in absolute quantification and partition-based detection. These improved quantitative precision and sensitivity in the detection of very low viral loads are particularly relevant for environmental surveillance and samples with low RNA concentrations. The biological relevance of low-positive dPCR signals was further supported by Sanger sequencing, which confirmed specific amplification of WNV even in samples close to the LoD.

The workflow for water was adapted from a patented method to improve recovery of insoluble fractions in environmental waters, where enveloped viruses may associate with suspended solids (Parkins et al., 2024; Ye et al., 2016). Experimental results supported improved recovery using the optimized protocol; however, the limited range of water types evaluated constitutes a limitation and extraction may vary with environmental conditions (turbidity, salinity, seasonal changes, etc.). Matrix heterogeneity and the presence of amplifications inhibitors represented important practical constraints. While increasing input volume for mosquito excreta improved sample representativeness without measurable inhibition, environmental waters required a reduced input volumes to accommodate the high variability in inhibitor content across wetlands and artificial breeding site waters. The systematic inclusion of internal controls, targeting both mosquito material and amplification inhibition, strengthens result interpretation in routine surveillance settings. Although dPCR is less sensitive to partial inhibition than qPCR–widely used in arboviruses surveillance (Daude et al., 2024; Mhamadi et al., 2023)– and allows absolute quantification without external standard curves, it still requires matrix-specific optimisations (Dingle et al., 2013; Guri et al., 2024). In addition, some robustness and repeatability parameters recommended by the dMIQE guidelines(dMIQE Group & Huggett, 2020) could not be fully addressed. These criteria were originally defined in the context of clinical and medical biology applications where homogenous samples and large volume of standardized material are typically available. The environmental matrices posed specific constraints; for instance, the strong heterogeneity in physicochemical properties across sites limited the relevance of conventional robustness testing based on a few representative matrices. We therefore prioritized the selection of a versatile protocol applicable across diverse environmental matrices. Beyond analytical performance, viral stability in environmental matrices is a critical factor for data interpretation, yet the persistence of viral particles and RNA in natural waters remains poorly characterised, a knowledge gap for sample preservation. Despite the large number of samples from 2024 and 2025 campaign and their diverse origins, the workflow proved robust and operationally applicable across highly heterogenous matrices. The precocity of detection relied in part on the use of digital PCR, which provides superior sensitivity and improved tolerance to inhibitors compared with qPCR. This advantage was illustrated by samples positive by dPCR but undetectable by qPCR, highlighting the added value of dPCR for identifying low-level viral RNA in environmental and mosquito-derived matrices. Future improvements in molecular workflows could enhance the sequencing potential of this matrix, including the use of targeted enrichment strategies, hybridization-based capture methods, or optimized multiplex amplification prior to sequencing. High-sensitivity sequencing technologies may enable the recovery of longer genomic fragments or even near-complete genomes directly from environmental waters. Such advances would provide confirmation of early detections but also enable genomic characterization of circulating WNV and USUV lineages from non-invasive environmental surveillance. Non-individual environmental surveillance proved its usefulness for early outbreak detection and surveillance in low-transmission areas(Cavany et al., 2022) and its potential as a complementary early-warning epidemiological tool alongside traditional and innovative surveillance systems(Zulli, Bowie, et al., 2025; Zulli, Zhang, et al., 2025).

The integrated use of MX and environmental water sampling provided a marked temporal advantage over conventional surveillance systems. WNV was identified in wetland water samples as early as the 1^st^ of July in 2024, approximately 4 weeks before the first alleged equine case(Grard et al., 2025), 7 weeks before the first detection in human blood(Grard et al., 2025), 40 days before the notification of the first autochthonous human case(Santé publique France, 2025), and 67 days before the first equine alert(RESPE, 2024a) in the region in 2024. In 2025, the first detection in water the 9^th^ of April preceded the first alert of autochthonous human case by 7 weeks, the first equine case in Hérault by 22 weeks, and in Gard by 17 weeks–whereas the departments of Aude and Pyrénées-Orientales have not issued any alerts at all. Such early detection underscores the capacity of environmental matrices to capture viral circulation before clinical or veterinary reported cases, a critical asset given the high proportion of asymptomatic WNV and USUV infections in humans and equines. Surveillance in local avian reservoirs and in migratory birds(Mancuso et al., 2022), although requiring substantial time and logistic investments, enables early detections–as early as March(Beaubaton et al., 2025; Münger et al., 2025). In comparison, water analysis detected the viruses from the beginning of the sampling campaign in April, indicating that our methodology may provide detection within comparable timeframe than traditional avian surveillance. However, for both target viruses, most detections in environmental water occurred weeks before detection in mosquito excreta, as highlighted by detections in both matrices within the same geographic area. For instance, in wetlands east of Montpellier, WNV was detected in water samples on the 6^th^ of august 2024, whereas mosquito excreta collected nearby tested positive only the 30^th^ of august 2024. Early detections in water samples likely reflect the capacity of the aquatic environments to integrate viral inputs from multiple potential sources such as excretion and decay of birds and mosquito or avian shedding (Nemeth et al., 2009), functioning as an early, integrative indicator of local and viral circulation and dynamics. For instance, more frequent WNV detection than USUV both years could reflect a higher WNV prevalence–hypothesis however challenged (Simonin et al, in prep–, competitive interactions–as WNV replicates faster and can outcompete USUV in co-infections, although prior USUV infection may reduce vector competence for WNV(Wang et al., 2020)– or greater viral stability or loads of WNV in the some matrices. Indeed, enveloped viruses like Orthoflaviviruses degrade rapidly in the environment, with rates affected by temperature, sunlight, salinity, water depth, weather, and habitat(Cox et al., 2025; Silverman & Boehm, 2021). Differences in prevalence between WNV and USUV in our samples could reflect their relative stability, as well as complex interactions within vectors and hosts, like larval infection through ingestion of viral particles(Hamel et al., 2024b) in circulation areas. Water-based screening can facilitates early localisation of transmission hotspots, while concomitant MX monitoring strengthens inference on vector involvement and enables viral sequencing and lineage identification for expanded genomic surveillance (Kittling & et al., 2026; Koch et al., 2024).

Although low-level viral RNA detection in environmental matrices cannot be interpreted as direct evidence of ongoing transmission or infectious virus, from a public health standpoint, this framework provides a robust basis for proactive surveillance to provide actionable risk signals when interpreted within a validated analytical framework and confirmed through repeated sampling and/or orthogonal confirmation. Indeed, beyond analytical sensitivity, the primary contribution of this multi-matrix strategy is its decision-support value at the territorial scale. Because WNV/USUV surveillance based on clinical and veterinary notifications is intrinsically delayed, early detection in environmental waters and mosquito excreta can be used to trigger time-sensitive actions, including vector control operations, communication to public and health professionals, intensified One-Health surveillance, and preparedness to arboviral expansion in traditionally and newly infected regions.(Abbas et al., 2025). Vector control targeting Culex mosquitoes remains essential for reducing WNV and USUV, both zoonotic viruses(Simonin, 2024), and transmission risks. Given Culex mosquitoes’ long-distance dispersal, short adult lifespan, and the dead-end role of humans and horses, adulticide measures are of limited relevance for vector control. (Vinogradova, 2000), larval control strategies therefore emphasize large-scale, regular larval habitat treatments, requiring early and accurate identification of viral circulation areas to optimize resource allocation(Comité de veille et d’anticipation des risques sanitaires (COVARS), 2024). A simplified yet diversified surveillance strategy as our methodology, covering broad geographic areas and sustained throughout the vector season, can enable such timely early warning signals and localisation to guide targeted control measures. Our methodology can be extended to urban vector control operation following confirmation of autochthonous human cases. The detection of WNV in the vicinity of Baillargues, north-east Montpellier, validated the capacity of these matrices to identify ongoing local viral circulation. Importantly, water samples enabled arbovirus detection in artificial larval habitats, including in areas of autochthonous dengue cases (such as Pérols in 2024), suggesting the potential for describing spatial and temporal co-circulations of WNV, USUV, and Aedes-borne viruses within shared environments.

In conclusion, our approach offers a scalable, field-compatible framework to support territorial decision-making (Fig. 8) by enabling earlier targeting of surveillance and vector control resources, and by improving preparedness for recurrent and expanding Culex-borne arboviruses. In practice, we propose an operational tiered interpretation of signals: (i) a single low-level detection above the LoB/LoD should prompt confirmatory re-sampling and local risk review, (ii) confirmed detections in repeated samples or in multiple matrices within the same area should trigger geographically targeted prevention measures, and (iii) sustained signals across weeks should support escalation of territorial preparedness and integrated One Health surveillance. The limitation of traditional systems is well illustrated in northern Italy, where some of the most structured WNV surveillance programs in Europe are implemented: over 60 traps and more than 100,000 mosquitoes collected annually achieve only a 2–5% WNV positivity rates during peak transmission(Barzon et al., 2022; Gobbo et al., 2025), while multi-million-euro blood donor screening programs detect a limited number of positive donations(Defilippo et al., 2022). These largely reactive and weakly predictive systems contrast with environmental surveillance, which provides an early warning of viral circulation (Defilippo et al., 2022; Paternoster et al., 2017), and allows preventive and anticipatory vector control operations. Multi-compartment surveillance, alongside predictive modelling and established One Health networks, outlines the next step for more anticipatory risk assessment and response capacity in endemic and emerging regions.

**Figure 8:**
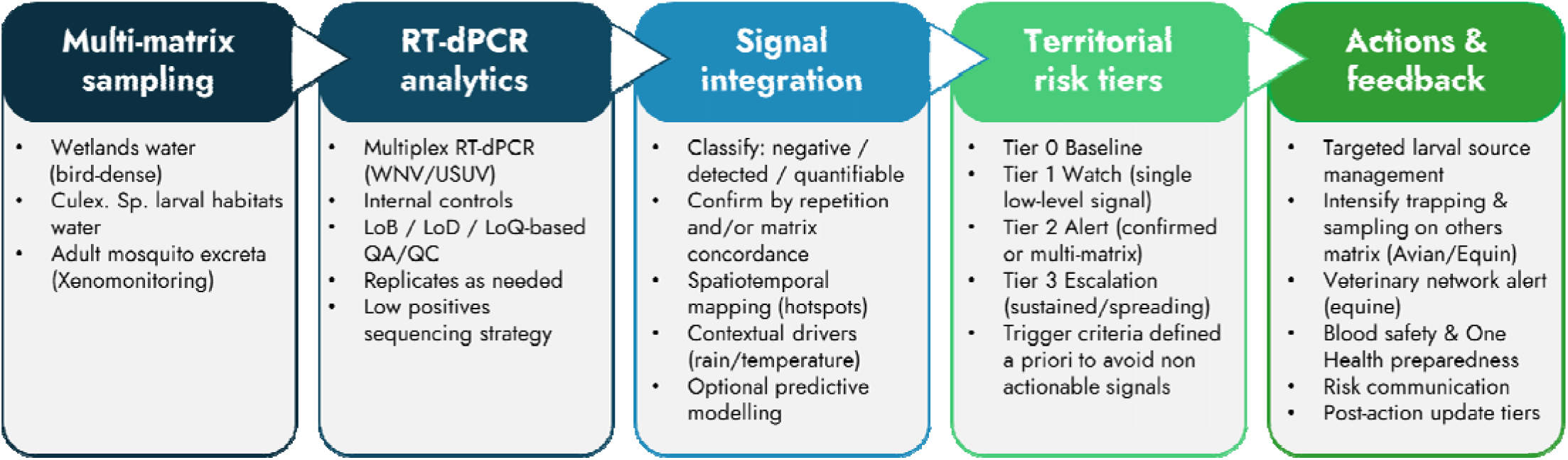
Territorial decision-support pipeline translating multi-matrix RT-dPCR surveillance signals into One Health actions.

## Acknowledgements

We gratefully acknowledge the financial support provided by La Région Occitanie, Bpifrance and France 2030, which made this study possible with regional I-DEMO project. We also thank Montpellier Méditerranée Métropole and MedVallée initiatives for their continuous institutional support. We further extend our gratitude to the environmental, conservation, healthcare and transport partners, including Montpellier Zoo, Écolothèque Montpellier 3M, ADENA, Pays de l’Or Agglomération, the Grand Site de France Camargue Gardoise, the LPO Centre de Sauvegarde, the Conservatoire du Littoral, CHU Toulouse, CHU Montpellier, for their logistical and technical assistance, which was essential for the implementation of field sampling and overall operational coordination. This work was carried out within the framework of the Camargue Health–Environment Zone Atelier (ZACAM), part of the Long-Term Socio-Ecological Research (LTSER) network, funded by the CNRS Institute of Ecology and Environment (CNRS-INEE). The consortium would also like to thank all current and former members of Altopictus, IAGE, and PCCEI who contributed to the creation and success of this project.

## Funding

This study was co-funded by the Région Occitanie, which covered 45% of the total project budget for IAGE and Altopictus, and the majority of the funds allocated to the University of Montpellier, with additional funding from the ExposUM Institute.

## Competing interests

All other authors declare no competing interests.

## Author Contributions

Conceptualization: GL, OC, AM, YS

Formal Analysis: JM, JR, KB, AL, CG, JH, AM

Funding Acquisition: GL, OC, AM, YS, FD

Investigation: JM, KB, RB, AL, JH, AF, GL, AM

Methodology: JM, JR, KB, RB, AL, CG, JH, GL, AF, OC, AM, YS

Project Administration and Supervision: GL, OC, AM, YS, FD

Visualization: JM, JR, CG, AM

Writing – Original Draft Preparation: JM, JR, KB, RB, AL, CG, JH, AM, YS

Writing – Review and Editing: JM, JR, KR, RB, AL, CG, GL, AF, OC, AM, YS

## Data Availability Statement

All experimental data will be made available from the Dryad Digital Repository before publication.

## Notes

### Competing Interest Statement

The authors have declared no competing interest.

