## Supplementary Figure 1 for "Early-warning surveillance of West Nile and Usutu viruses in water and mosquito excreta using digital PCR"

**Supplementary figure S1: Example of positive and negative raw results for WNV (A) and USUV (B).**

**Supplementary Table S1: Details of sampling points for 2024 and 2025 campaign.** USUV and WNV detection indicate the detection at least once of the virus at the sampling point.

Excel file: SuppS1_List_Sites.xlsx

**Supplementary Table S2. Primers and probe targeting WNV and USUV viruses**

**Supplementary Table S3: dMIQE2020 checklist for authors, reviewers and editors.**

Excel file: SuppS3_dMIQE2020.xlsx

**Supplementary Table S4. Stability test Pan-mosquito and WNV viruses over time**

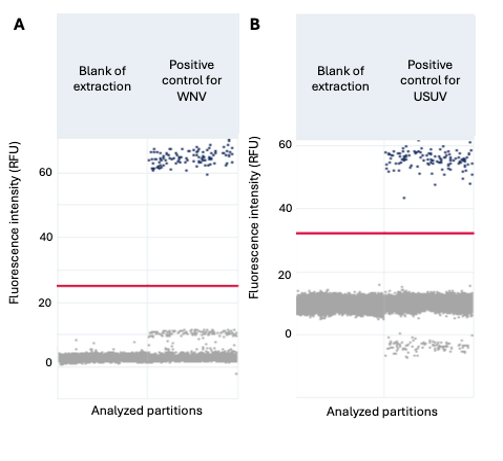

**Supplementary figure S1: Example of positive and negative raw results for WNV (A) and USUV (B).** Blue dots represent positive partitions, grey dots negative partitions. The red line represent the threshold separating positive and negative partitions.

| **Supplementary Table S2. Primers and probe targeting WNV and USUV viruses** | | | | |
| --- | --- | --- | --- | --- |
| Target | Oligonucleotids | Amplicon size (bp) | Sequence accession number | Reference |
| WNV | F 5' CTGTGTGAGCTGACAAACTTA 3' | 105 | Lineage 1 NC_009942.1 Lineage 2 NC_001563.2 | (Kolodziejek et al., 2014) |
|  | P 5' TGCGAGCTGTTTCTTAGCACGA 3' |  |  |  |
|  | R 5' CTCCTGGTTTCTTAGACATCG 3' |  |  |  |
| USUV | F 5' CATCGTTCTCGACTTTGACT 3' | 88 | NC_006551 | (Jöst et al., 2011) |
|  | P 5' ACCGTCACAATCACTGAAGCAT 3' |  |  |  |
|  | R 5' AGTAGTTCTTATGGAGGGTCC 3' |  |  |  |

| **Supplementary Table S4. Stability test Pan-mosquito and WNV viruses over time** | | | | | | |
| --- | --- | --- | --- | --- | --- | --- |
| Sample | Matrix | Municipality | Sampling date | Shelf life at 4°C(days) | [Pan-mosquito] (Copies/L) | [WNV] (Copies.L) |
| 1 | Water - Artificial breeding site | Baillargues, France | 09/09/2024 | 0 | 1,62E+10 | 1,25E+05 |
| 1 | Water - Artificial breeding site | Baillargues, France | 09/09/2024 | 8 | 1,48E+10 | 2,47E+05 |
| 1 | Water - Artificial breeding site | Baillargues, France | 09/09/2024 | 15 | 1,33E+08 | 3,56E+05 |
| 1 | Water - Artificial breeding site | Baillargues, France | 09/09/2024 | 23 | 6,12E+06 | 1,09E+05 |
| 1 | Water - Artificial breeding site | Baillargues, France | 09/09/2024 | 25 | 2,40E+09 | 5,52E+04 |
| 1 | Water - Artificial breeding site | Baillargues, France | 09/09/2024 | 28 | 1,76E+06 | ND |
| 1 | Water - Artificial breeding site | Baillargues, France | 09/09/2024 | 42 | 1,40E+06 | ND |
| 2 | Water - Wetlands | Baillargues, France | 09/09/2024 | 0 | 1,46E+10 | 6,38E+04 |
| 2 | Water - Wetlands | Baillargues, France | 09/09/2024 | 8 | 7,28E+09 | 5,63E+04 |
| 2 | Water - Wetlands | Baillargues, France | 09/09/2024 | 15 | 4,57E+09 | ND |
| 2 | Water - Wetlands | Baillargues, France | 09/09/2024 | 15 | 4,06E+09 | 1,12E+05 |
| 2 | Water - Wetlands | Baillargues, France | 09/09/2024 | 23 | 1,31E+08 | 1,10E+05 |
| 2 | Water - Wetlands | Baillargues, France | 09/09/2024 | 25 | 5,79E+07 | 1,12E+05 |
| 2 | Water - Wetlands | Baillargues, France | 09/09/2024 | 28 | 4,01E+07 | 5,83E+04 |
| 2 | Water - Wetlands | Baillargues, France | 09/09/2024 | 42 | 2,52E+07 | ND |

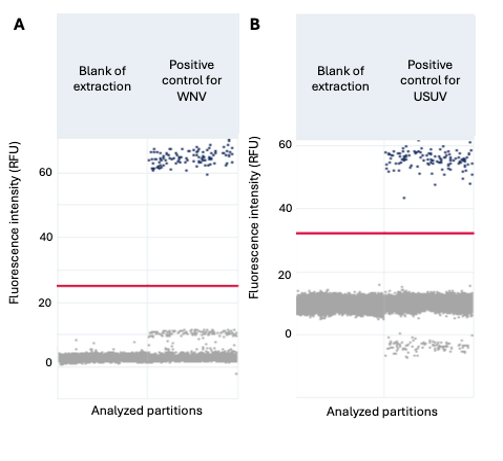

**Figure S1. Example of positive and negative raw results for WNV (A) and USUV (B)**. Blue dots represent positive partitions, grey dots negative partitions. The red line represent the threshold separating positive and negative partitions.
